# *In vitro* evaluation of immune responses to bacterial hydrogels for the development of living therapeutic materials

**DOI:** 10.1101/2022.09.16.508081

**Authors:** Archana Yanamandra, Shardul Bhusari, Aránzazu del Campo, Shrikrishnan Sankaran, Bin Qu

## Abstract

In living therapeutic materials, organisms genetically programmed to produce and deliver drugs are encapsulated in porous matrices or hydrogels acting as physical barriers between the therapeutic organisms and the host cells. The therapeutic potential of such constructs has been highlighted in *in vitro* studies, but the translation to *in vivo* scenarios requires evaluation of the immune response to the presence of the encapsulated, living organisms. In this study, we investigate the responses of human peripheral blood mononuclear cells (PBMCs) exposed to a living therapeutic material consisting of engineered *E. coli* encapsulated in Pluronic F127-based hydrogels. The release of inflammation-related cytokines (IL-2, IL-4, IL-6, IL-10, IL-17A, TNFα and IFNγ) and cytotoxic proteins (granzyme A, granzyme B, perforin, granulysin, sFas, and sFasL) in response to the bacterial hydrogels, as well as the subsets of natural killer cells and T cells after exposure to the bacterial hydrogel for up to three days were examined. In direct contact with PBMCs, both *E. coli* and its endotoxin-free variant, *ClearColi*, induce apoptosis of the immune cells and trigger IL-6 release from the surviving cells. However, we found that encapsulation of the bacteria in Pluronic F127 diacrylate hydrogels considerably lowers their immunogenicity and practically abolishes apoptosis triggered by *ClearColi*. In comparison with *E. coli*, free and hydrogel-encapsulated *ClearColi* induced significantly lower levels of NK cell differentiation into the more cytolytic CD16^dim^ subset. Our results demonstrate that *ClearColi*-encapsulated hydrogels generate low immunogenic response and are suitable candidates for the development of living therapeutic materials for *in vivo* testing to assess a potential clinical use. Nevertheless, we also observed a stronger immune response in pro-inflammatory PBMCs, possibly from donors with underlying infections. This suggests that including anti-inflammatory measures in living therapeutic material designs could be beneficial for such recipients.

## Introduction

Living therapeutic materials (LTMs) combine live cells with non-living materials to create devices with the potential to deliver therapeutic molecules in response to patient’s needs^1,2^. Biosensors, wound patches, tissue adhesives and drug delivery implants are reported examples that demonstrate the interest of this technology for different biomedical scenarios. Bacteria are frequently used as living component in LTMs due to their simple and adaptable nutritional requirements, and their robustness to survive harsh conditions (e.g. pH, hypoxia, osmosis) or processing steps for storage (e.g. freeze drying). These living cells are encapsulated in polymeric matrices, typically agarose ^3, 4, 5^, alginate ^6^ or Pluronic F127 ^7, 8, 9^. At adequate composition, the encapsulating matrices allow diffusion of nutrients, gas, and metabolites and maintain the viability and function of the organisms while physically separating them from antagonistic external entities like competitive microbes or immune cells.

For the use of LTMs in biomedical applications, one major/decisive aspect is the immune responses they might elicit. Immunogenicity of LTMs can be associated to the material and to the living bacterial components. It is important to highlight that in LTMs, bacteria are not directly exposed to the host cells and, therefore, possible immunogenicity should be associated with secreted substances or cell debris or with the properties of the encapsulating hydrogel, but not with a direct bacteria-host cell contact. The inflammatory host reactions are complex and involve neutrophils as the first dominant responder, followed by macrophages ^10^. CD4^+^ T helper cells and immune killer cells (CD8^+^ cytotoxic T lymphocytes and natural killer cells) also participate in inflammation responses ^11^. Upon detection of foreign bodies, inflammation-related cytokines (e.g. IL-6, IL-8, IL-10, tumor necrosis factor α (TNFα), interferon-gamma (IFNγ), and IL-17A) and cytotoxic proteins (e.g. perforin, granzyme A, granzyme B, granulysin, soluble Fas and soluble Fas ligand) are released by the effector immune cells ^12, 13, 14, 15, 16, 17, 18, 19, 20, 21^. Profiles of these cytokines and cytotoxic proteins can be therefore used as markers to evaluate LTM immunogenicity. Certain immune cells, including T cells and NK cells, can differentiate into functionally different subsets upon exposure to external stimuli ^22, 23^, offering another aspect to access LTM-induced immune response.

In this paper, we examine how the immune cells react to LTMs made from a commonly-reported material-bacteria combination: Pluronic F127-based hydrogels and *E. coli* strains. Pluronic F127 is a synthetic polymer approved for medical use and frequently used in drug delivery formulations. Pluronic solutions at concentrations >5 wt% and temperatures above 14°C self-assemble into micelles to form associative (physical) hydrogels ^24, 25^. This thermoreversibility facilitates the use of Pluronic to encapsulate drugs or cells. *E. coli* has been frequently used in LTM design due to the availability of a broad genetic toolbox and medically relevant strains (e.g. Nissle 1917, *ClearColi*) ^1^. *E. coli* hosts lipopolysaccharides (LPS) on the outer membrane, that engage with Toll-like receptor 4 (TLR4) and trigger strong endotoxic responses, i.e. pro-inflammatory immune responses in mammals especially humans, which can result in a life-threatening cytokine storm. In the endotoxin-free strain *ClearColi* BL21DE3, the 7 genes that encode for LPS synthesis are deleted. Lysates of this strain have been proven to not elicit endotoxic responses ^26^. In our recent work, we described light-responsive LTMs containing *ClearColi* engineered to produce and secrete a fluorescent protein as a model protein and deoxyviolacein as a potential antimicrobial drug ^3, 4^. We demonstrated that drug profiles could be tuned by light irradiation parameters and that release could be sustained for over a month in both cases, suggesting that such LTMs could be used to develop *in situ* controllable drug delivery devices. Inspired by these positive results, and with the aim to extend our future studies to *in vivo* scenarios, we examine here how donor-derived peripheral blood mononuclear cells (PBMCs) react to LTMs made of Pluronic F127 based hydrogels containing *ClearColi*. For comparison, similar studies with *E. coli* BL21DE3 (endotoxic) and empty Pluronic F127 gels were also performed. The results of our study suggest that LTMs containing (and retaining) the endotoxin-free *ClearColi* within Pluronic hydrogels do not cause strong immune reactions by PBMCs, indicating the potential suitability of this combination for LTM development. We also noticed that donors with pro-inflammatory status (high spontaneous IL-2 release) did show inflammatory response to the LTMs and, therefore, preventive strategies might need to be considered for the applicability of LTMs to such patients.

## Materials and Methods

### Antibodies and reagents

The following antibodies were purchased from BioLegend: BV421 anti-human CD3 antibody, PerCP anti-human CD8 antibody, PE anti-human CD45RO antibody, Alexa647 anti-human CCR7 antibody, APC anti-human CD56 antibody, PerCP anti-human CD16 antibody. Legendplex human CD8/NK panel was also purchased from BioLegend. Lymphocyte separation medium 1077 was purchased from PromoCell.

### Bacteria cultures

*E. coli* BL21(DE3), from NEB (C2527H), and its endotoxin-free variant, *ClearColi* BL21(DE3), from Lucigen (60810-1), were used in this study. They were transformed with the plasmid pUC19 to provide them with ampicillin resistance and minimize the risk of contamination in the culture. Transformation in *E. coli* was performed by heat-shock and in *ClearColi* by electroporation as described by the manufacturer of these competent cells. Bacterial cultures were grown for 16 h at 35°C, 180 rpm in LB Miller medium supplemented with 50 µg/mL of ampicillin to an optical density at 600 nm wavelength (OD600) between 0.5 – 1.

### Functionalization of glass coverslips with (3-acryloxypropyl)trimethoxysilane

13 mm glass coverslips were sonicated in ethanol for 5 min and then rinsed with ethanol. The coverslips were treated with 1% v/v solution of (3-acryloxypropyl)trimethoxysilane (APS, Merck - Sigma Aldrich) in ethanol for 1 h. The coverslips were then rinsed in ethanol and dried for further use.

### Preparation of bacteria-encapsulated thin-film hydrogel constructs

Pluronic F127 (Plu, MW~12600 g/mol, Sigma-Aldrich) and Pluronic F127 diacrylate (PluDA) ^27^ with substitution degree of 70% was used for the studies. Stock solutions of 30% (w/v) Plu and PluDA in milliQ water with Irgacure 2959 photoinitiator at 0.2% w/v were prepared and stored at 4°C. 1:1 mixtures of the stock solutions of Plu and PluDA at 4°C were added to the bacterial suspension (OD600 of 0.5, ~ 4 × 10^7^ cells/mL) at 9/1 (v/v) ratio to achieve a final OD600 of 0.05. The solutions were stored on ice to keep the Plu/PluDA solutions in liquid form. The Plu/PluDA/bacteria mixture was vortexed for 30 sec to obtain a homogeneous dispersion. 2 µL of the Plu/PluDA/bacteria mixture was placed on coverslips previously functionalized with (3-acryloxypropyl)trimethoxysilane and left at room temperature for 10 min to form the physical hydrogel (Supporting information Figure S1). The silanization step ensures covalent bonding of the PluDA component with the glass substrate during the upcoming photopolymerization step.

The bacteria/Plu/PluDA physical hydrogel was covered by a PluDA envelope to avoid bacteria outgrowth. For this purpose, 30 µL of PluDA solution was dropped on parafilm and allowed to physically crosslink at room temperature for 10 min. The coverslip with the bacterial gel was then placed on top of the gel on the parafilm. The hydrogel sandwich was exposed to UV light (365 nm, 6 mW/cm^2^) using a OmniCure Series 1500 lamp for 60 sec. After additional 10 min, the parafilm was peeled off.

The thin-film construct was incubated at 37°C with 5% CO_2_ for 3 days before incubation with the immune cells. In this time bacteria grew into colonies inside the Plu/PluDA hydrogel core layer. Samples showing bacteria leakage (~1 out 10) were not used for the study. From our and others’ previous studies,^7, 9, 28^ we consider that the Pluronic F127 based gels allow free diffusion of nutrients. Plu/PluDA hydrogels might leak non-crosslinked chains to the medium during the first hours of incubation.

### Live/Dead assay

The LIVE/DEAD BacLight Bacterial Viability kit (Thermo Fisher Scientific L7012) was used. Stock solutions of the stain were stored at −20°C. To make the working solution, 3 µL of the stock solutions were added to 1 mL of phosphate saline buffer (PBS). The bacterial hydrogels were washed with PBS to remove traces of the RPMI medium and incubated with 100 µL of the stain solution for 20 min. The samples were washed with 0.5 mL PBS once and imaged under the microscope. For fluorescence imaging, filter channels BZ-X Filter GFP OP-87763 (excitation 470/40, emission 525/50) and BZ-X Filter OP-87764 (excitation 545/25, emission 605/70) and a 4× objective (Keyence Plan Apochromat, numerical aperture 0.20, working distance 20 mm) were used. Image processing and analyses were performed using Fiji edition of ImageJ (ImageJ Java 1.8.0). The thin-films were also imaged by Zeiss LSM 880 confocal laser scanning microscopy (CLSM) at days 3 and 7. The exposure conditions were optimized for minimizing cell photodamage using the objective LD C-Apochromat 40×/1.1 W Korr M27, detection wavelength 493-584 nm and 584-718 nm, laser wavelength of 488 and 583 nm and power of 0.2 and 1.5 % respectively for live and dead bacterial populations. Z-stacks of 100.204 µm were taken in a z-step size of 0.65 µm. Images of a size of (xy) 283.77 × 283.77 µm were acquired (512 × 512 pixels), two-fold line averaging, and pixel dwell time of 1.52 µs. The digital gain value was 760 and 720 for live and dead bacterial populations. Imaris software (Version 9.0, Bitplane) was used to process CLSM image z-stacks to create three‐dimensional images.

### ATP assay

Bacterial cell viability was assessed by the CellTiter-Glo luminescent cell viability assay (Promega) that quantifies ATP released in the culture medium as an indicator of metabolically active cells. A 1:1 mixture (25 μL each) of the CellTiter-Glo reagent and the supernatant was shaken for 5 min and the luminescence was measured using a Tecan Infinite M200-Pro plate reader. The experiments were performed for supernatants from different culture times.

### Protein/DNA release studies

To quantify the amount of protein released in the supernatant medium of the thin films, a commercial BCA protein assay (Pierce BCA Protein Assay Kit, Thermo Fisher Scientific) was used. Quantification by absorbance measurements at 562 nm was performed using a Tecan Infinite 200 PRO microplate reader. To check the release of DNA into the supernatant, agarose gel electrophoresis was performed with the cell free supernatants mixed with the Tritrack Gel Loading Dye in a 3:1 ratio and loaded on the 1.5% (w/v) agarose gel premixed with SyBr Safe gel staining dye. The samples were run at constant voltage (100V) and visualized in the ChemiDoc TransIlluminator under UV light irradiation.

### Isolation and cell culture of PBMCs

Human peripheral blood mononuclear cells (PBMCs) were isolated from Leucocyte reduction system chambers of healthy donors employing gradient centrifugation method where Lymphocyte Separation Medium 1077 was used. Virological test excluded infection by HIV and hepatitis B in the donors. Research carried out for this study is authorized by the local ethic committee (declaration from 16.4.2015 (84/15; Prof. Dr. Rettig-Stürmer)).

To test the immune response to LTMs, PBMCs were cultured at 37°C with 5% CO_2_ for 3 days in RPMI-1640 medium (Thermo Fisher Scientific) supplemented 10% FCS (Thermo Fisher Scientific) and 1 % ampicillin at a cell density of 3×10^6^/mL per condition. Eight different conditions were used in the experiments: PBMCs alone (Ctrl), blank gel (without bacteria), *ClearColi* encapsulated gel, *E. coli* encapsulated gel, *ClearColi* in a transwell insert, *E. coli* in a transwell insert, *ClearColi* in direct contact with PBMCs, *E. coli* in direct contact with PBMCs. Bacteria suspensions with initial OD 0.05 were used for the studies.

### Identification of subpopulations and subsets

PBMCs were collected on day 3 and washed twice with PBS/0.5% BSA, followed by staining with corresponding antibodies as specified in the figure notations for 30 min at 4°C in dark.

Control IgG was used to gate the positive cells. FACSVerse flow cytometer (BD Biosciences) was used for data acquisition and FlowJo v10 (FLOWOJO, LLC) for data analysis.

### Multiplex cytokine assay

100 μL of cell culture media was collected carefully on days 1, 2 and 3 from each condition, centrifuged at 1000 g for 10 min. Supernatant was collected and stored at −80°C until further analysis. Legendplex human CD8/NK panel (BioLegend) was used to determine cytokine profiles of the samples according to the manufacturer’s instructions. Culture media was used as the blank control.

### Statistical analysis

GraphPad Prism 9 Software (San Diego, CA, USA) was used for statistical analysis. The differences between two columns were analyzed by the ratio paired t-test.

## Results

In our previous studies we demonstrated that *ClearColi* encapsulated in 30 wt% Plu/PluDA hydrogels were able to grow and keep their metabolic function ^7, 9^. The growth and activity of the encapsulated bacteria was dependent on the PluDA ratio in the hydrogel, which controls the covalent crosslinking degree of the network. In PluDA hydrogels, bacteria showed the lowest growth and remained confined within the hydrogel over days. In 1:1 Plu/PluDA hydrogels bacteria showed the highest protein production rate but they were able to grow out of the gels. For the current studies, we processed hydrogels in a bilayer thin film format, where 1:1 Plu/PluDA hydrogels containing bacteria were enveloped by PluDA hydrogels without bacteria (Figure 1A, Supplementary information Figure S1). This bilayer format supported bacterial growth and metabolic production while it prevented outgrowth and escape of the bacteria during at least 7 days. It also facilitated microscopy imaging of the encapsulated bacteria. At the used seeding density, bacteria grew inside the thin film hydrogel and formed individual colonies, as previously reported. The bacteria within these colonies remained viable for at least 7 days according to the live/dead staining (Figure 1B). The metabolic activity of the encapsulated bacteria was assessed by quantifying the extracellular ATP. This assay allows detection of the metabolic activity of bacterial cultures at concentrations <50 nM ^29^. In a classical bacteria culture, extracellular ATP peaks during the early log phase and decreases in the stationary phase ^29^. A similar bell-shaped curve was observed in the supernatant of bilayer hydrogels. After a 6 h lag period, nearly 0.15 nM ATP was detected for the following 22 hours, followed by a progressive drop (Supplementary information, Figure S2). We infer that the bacteria within the first 6 to 24 h of encapsulation and incubation within medium, are in the log phase, whereas bacteria at 48 and 72 h have reached the stationary phase. This time scale correlates with the observed colony growth rates discussed previously.

**Figure 1:**
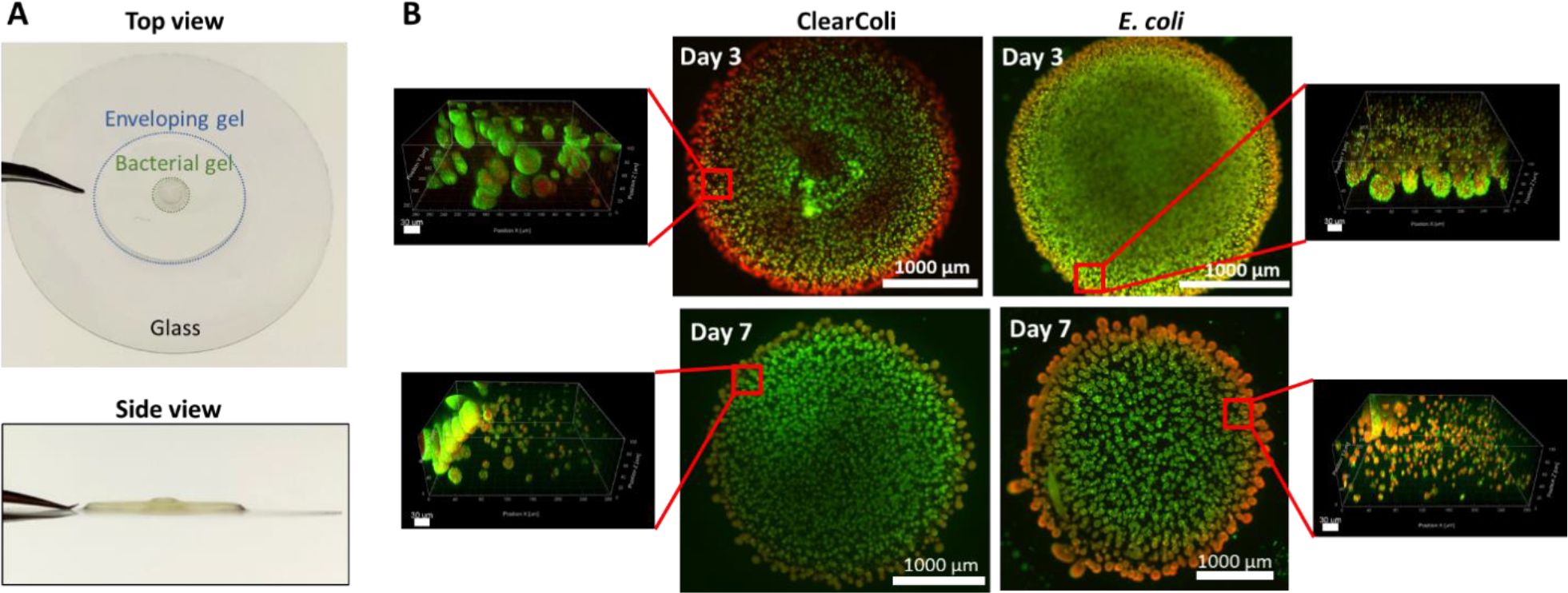
Bacteria/hydrogel thin-film constructs. **A)** Thin films (top and side view) showing the enveloping and bacterial Plu/PluDA gels. **B)** Confocal and fluorescence microscopy images of bacteria colonies of *ClearColi* and *E. coli* embedded in the hydrogels after staining with live-dead assay. The images show the presence of a viable population at day 3 and 7 post fabrication. (green/SYTO9 = live, red/propidium iodide = dead).

Analysis of the incubation medium after 4, 5, and 6 days of incubation with BCA protein assay and gel electrophoresis did not reveal significant release of proteins or DNA from the gels (Supporting information Figure S3) ^31, 32^. This result suggests that no notable cell lysis occurs within this time.

### PBMC response to ClearColi, E. coli and PluDA hydrogels

We co-cultured PBMCs with the bacterial hydrogels and analyzed PBMC-released cytokines and cytotoxic proteins by multiplex cytokine assays for up to 3 days, since this is the relevant timespan for T cells to be fully activated ^33^. Note that the bacterial hydrogels were kept in culture for 3 days before they were cocultured with the PBMCs. This step ensured that the bacteria had grown to their maximum colony sizes in all samples and allowed us to verify that they did not grow or leak out from the thin films. PBMCs from healthy donors were used as they contain innate immune cells (e.g. monocytes and NK cells) as well as adaptive immune cells (e.g. T cells and B cells) ^34^ (Supplementary information Figure S4).

As control experiments, we quantified the immunogenicity of *E. coli* and *ClearColi* suspensions in direct contact with PBMCs, or physically separated by a nano-porous membrane in a transwell plate, i.e. ensuring exchange of bacteria-produced soluble factors (Figure 2A). Profiles of PBMC-released cytokines and cytotoxic proteins were analyzed on days 1, 2 and 3. The cytokines IL-2 ^35, 36^, IL-4 ^37^, IL10 ^38^, IL-6 ^39, 40, 41^, IL-17A ^42^, TNFα ^43^ and IFNγ ^44^ were selected for the panel as they play essential roles in both innate and adaptive immune response against infection. In addition, cytotoxic proteins (sFas, sFasL, GzmA, GzmB, perforin and granulysin) were included in the panel to access reaction of immune killer cells, mainly NK cells and CTLs, to bacteria-encapsulated hydrogels. When PBMCs were in direct contact with *E. coli*, IL-6 was predominantly released. TNFα and IFNγ were also detectable but at much lower levels (Figure 2B, left panel). *ClearColi* elicited release of the same cytokines, but at half the level of *E. coli* (Figure 2B, left panel). When bacteria and PBMCs were separated by the transwell membrane with 400 nm pores, IL-6 release was triggered from day 2 onwards, which is one day later compared to the direct contact condition (Figure 2B). This result indicates that soluble factors from the bacteria, and not only direct contact, can trigger cytokine release. In the transwell condition, compared to *ClearColi* IL-6 release was 2-fold higher for *E. coli* (Figure 2B, right panel). These results confirm that the lack of endotoxic LPS in *ClearColi* reduces immunogenicity, no matter if there is direct contact between bacteria and host or not. Immune cells, especially NK cells, CD4^+^ T cells and CD8^+^ T cells, can differentiate into functionally different subsets upon activation. Thus, we also analyzed differentiation of these cells in response to the blank hydrogels on day 3. It should be noted that more than half of PBMCs were apoptotic both in the direct bacterial contact (Figure 2C) and in transwell-separated (Figure 2D) cases, as identified by positive PI-staining. In comparison, for the controls, i.e. no bacteria present, the apoptotic fractions were lower than 3% in both cases (Figure 2C, D). In the remaining viable PBMCs, NK cells were polarized to the CD16^dim^ subset by *E. coli* but not by *ClearColi* (Figure 2E, F). In terms of CD4^+^ and CD8^+^ T cells, four subsets were examined: naive (CD45RO^−^CCR7^+^), central memory (CM) (CD45RO^+^CCR7^+^), effector memory (EM) (CD45RO^+^CCR7^−^) and effector (CD45RO^−^CCR7^−^) cells. No change in differentiation of CD4^+^ and CD8^+^ T cells was detected (Figure 2G-J), although due to bacteria-induced apoptosis, the number of detected events was reduced as shown by the shrunk sizes of subsets compared to the condition without bacteria (Figure2 G, I).

**Figure 2:**
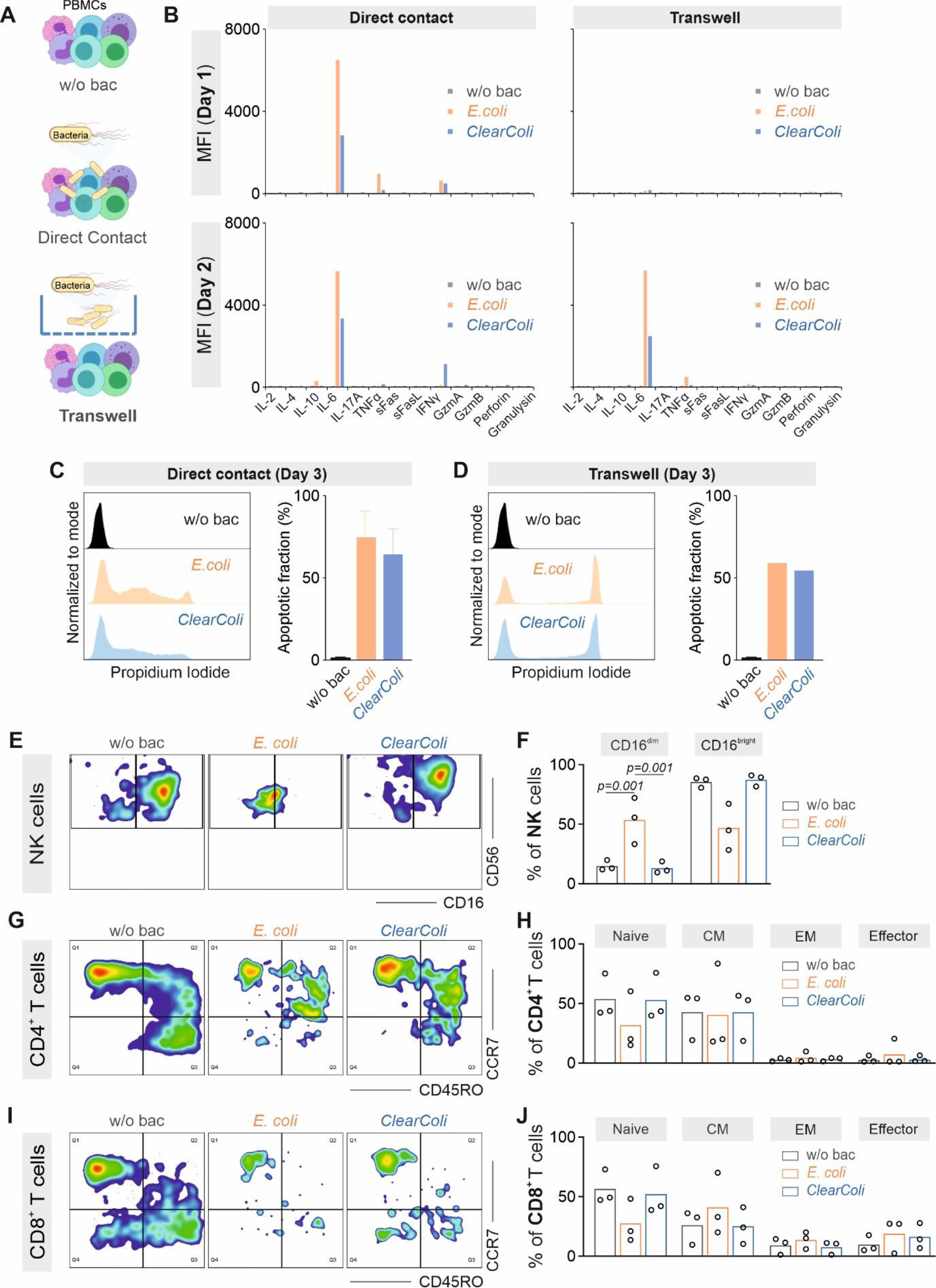
Immune response of PBMCs exposed to bacteria. **A)** Schematic for the different conditions tested: PBMCs cultured without bacteria (w/o bac), with bacteria in the same well (Direct Contact), or with bacteria in a transwell, where PBMCs and bacteria are separated by a porous membrane that precludes direct contact (Transwell). **B)** Profiles of cytokines and cytotoxic proteins released by PBMCs determined by the multiplex cytokine assay. PBMCs were cultured with or without bacteria in direct contact or transwell as indicated in the figure. The supernatant was collected at the indicated time points. Results were obtained from three donors and one representative donor is shown here. **C-D)** Viability of PBMCs on day 3 after direct contact with bacteria (**C**) or exposed to the soluble factors in a transwell (**D**). On day 3, PBMCs were stained with propidium iodide (PI) and PI positive PBMCs (higher PI intensity value on x-axis) were considered as apoptotic/dead cells. Results were from three donors in (**C**) and one representative donor in (**D**) is shown. **E-J**) Differentiation of NK cells (**E, F**), CD4^+^ T cells (**G, H**) and CD8^+^ T cells (**I, J**) examined using the indicated surface markers. Results were from three donors. CM: central memory cells. EM: effector memory cells.

Next, we examined the response of the PBMCs to the hydrogels without bacteria. The levels of cytokine or cytotoxic protein release were comparable to those of PBMCs controls and negligible compared to PBMCs in direct contact with *E. coli* (Figure 3).

**Figure 3:**
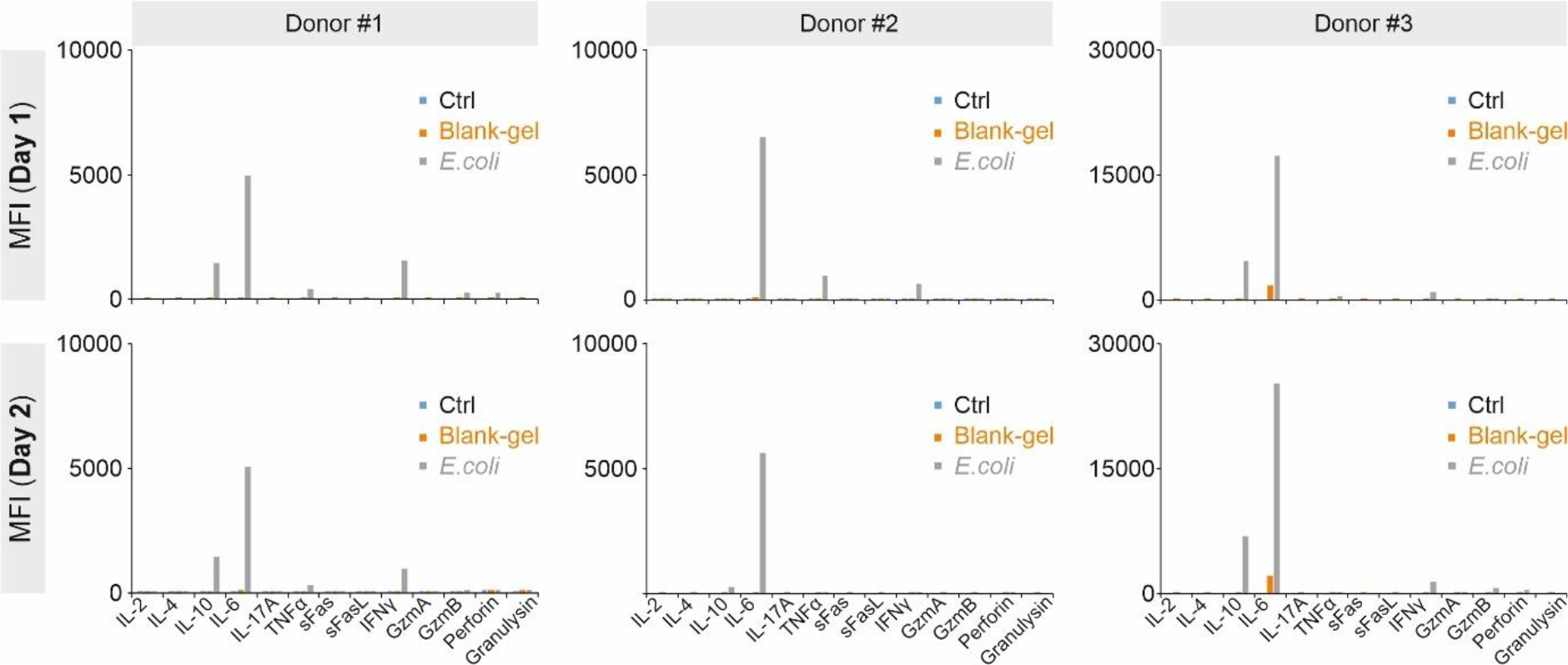
Profiles of cytokines and cytotoxic proteins in response to empty hydrogels. PBMCs were cultured without bacteria or gels (Ctrl), with empty hydrogels (Blank-gel), or in direct contact with *E.coli* (*E.coli*). Profiles of cytokines and cytotoxic proteins released by PBMCs were determined by the multiplex cytokine assay. The supernatant was collected at day 1 or day 2. Results were from three donors. MFI: mean fluoresce intensity.

NK cell differentiation analysis on day 3 revealed no differences in the fraction of the more cytolytic CD56^+^CD16^dim^ subset and the less cytolytic CD56^+^CD16^bright^ subset ^45^ between controls and samples co-incubated with the blank hydrogels (Figures 4A, B). Similarly, the presence of blank hydrogels did not alter the fraction of the CD4^+^ and CD8^+^ T cell subsets (Figures 4C-F). Taken together, these results indicate that the thin-film blank hydrogel construct is inert to immune cells *in vitro* and, therefore, Pluronic hydrogels are a reasonable material choice for the development of LTMs.

**Figure 4:**
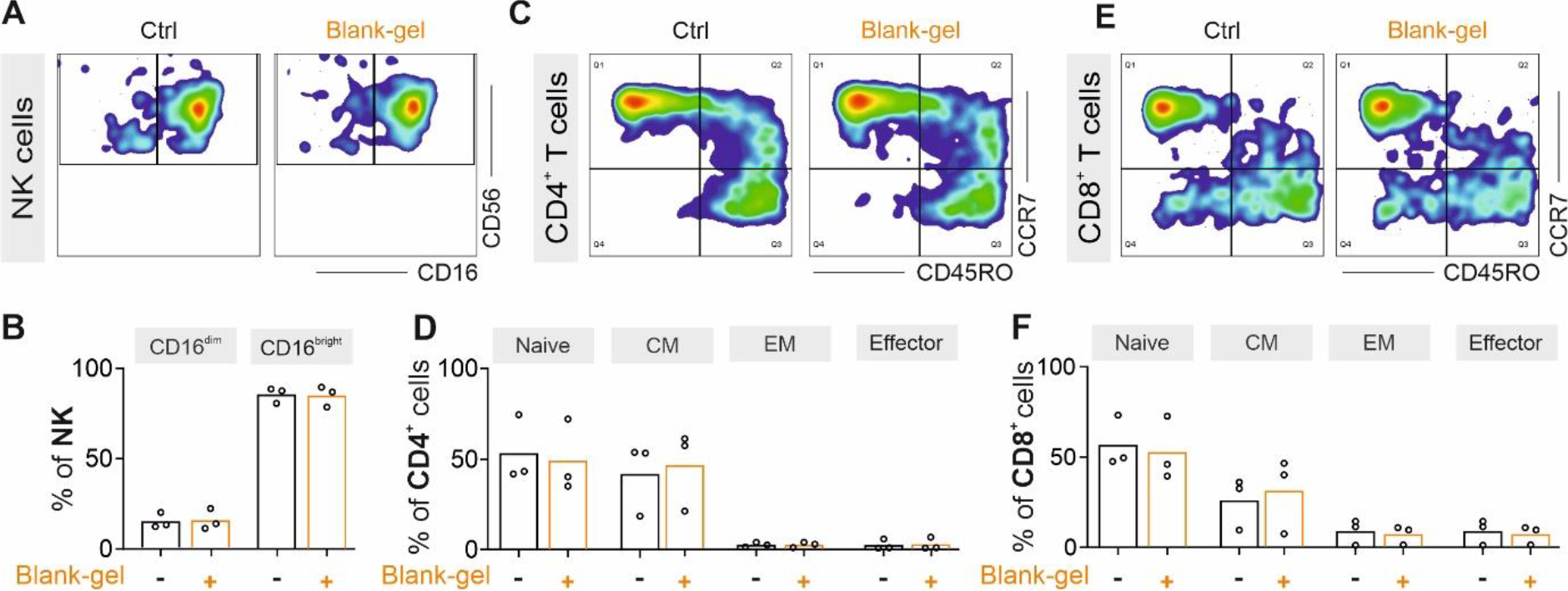
Differentiation of NK and T cells in response to empty hydrogels. PBMCs were cultured without gels (Ctrl) or with empty hydrogels (Blank-gel) for three days. Differentiation of NK cells (**A, B**), CD4^+^ T cells (**C, D**) and CD8^+^ T cells (**E, F**) were examined using indicated surface markers. Results were from three donors. CM: central memory cells. EM: effector memory cells.

### Characterization of immune responses to encapsulated E. coli and ClearColi hydrogels

We analyzed the response elicited by the bacterial thin-film hydrogels. PBMCs were co-cultured with the bacterial gels for 3 days and the viability of PBMCs was examined. In the case of *E. coli* hydrogels, 25 – 50% of the immune cells were found to be apoptotic on day 3, whereas for *ClearColi*-gels, the apoptotic fraction was reduced to 5% (Figure 5A). Compared to the conditions with direct contact to *ClearColi*, encapsulated bacteria in Pluronic hydrogel retain viability of immune cells at > 90% levels. This is a significantly higher viability than in the experiments where *ClearColi* were in direct contact or physically separated from the host cells through a porous membrane (<40 %).

**Figure 5:**
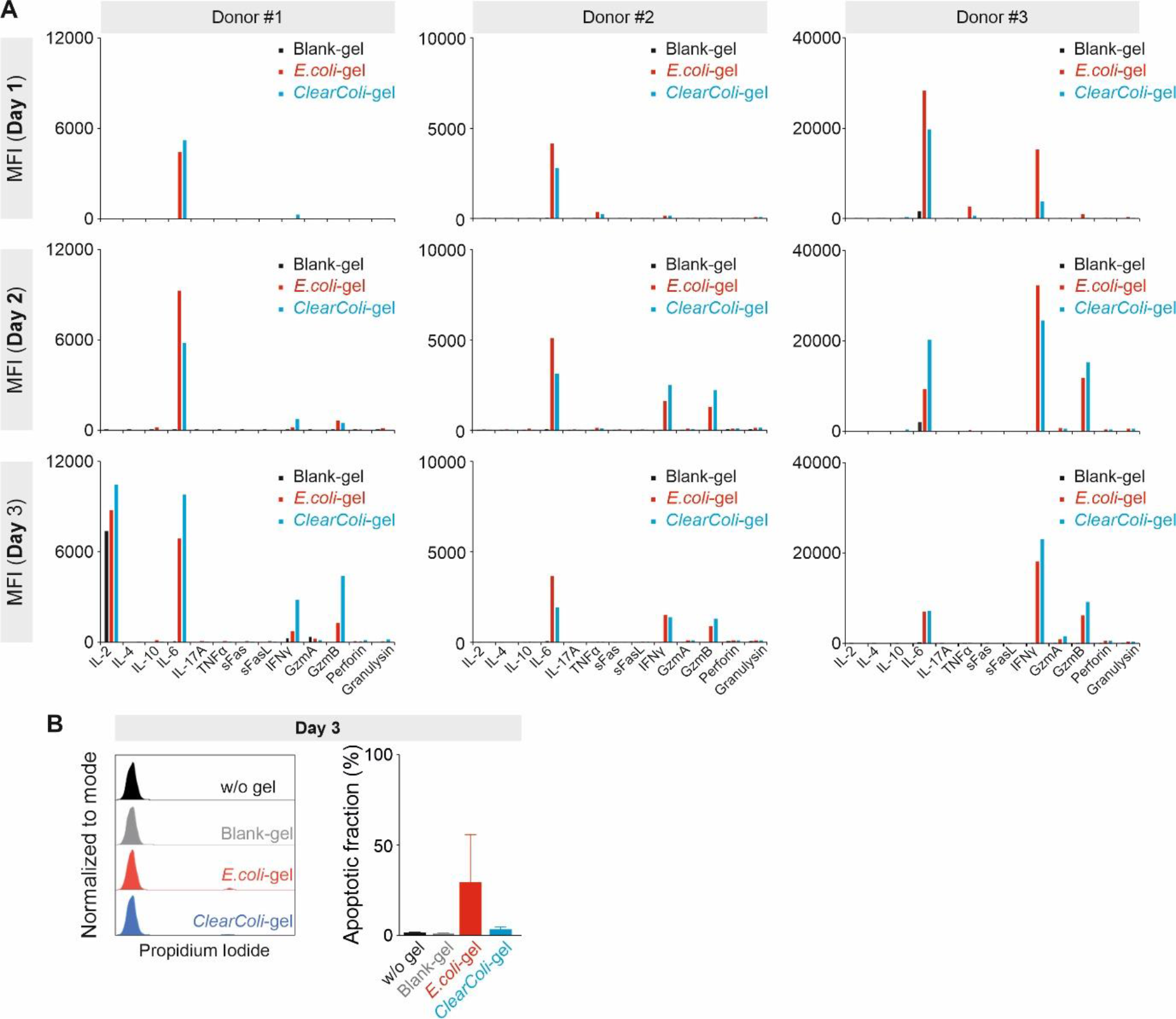
Profiles of cytokines and cytotoxic proteins in response to bacteria encapsulated hydrogels. PBMCs were cultured with empty hydrogels (Blank-gel), *E. coli*-encapsulated (*E.coli*-gel) or *ClearColi*-encapsulated gels (*ClearColi*-gel). **A)** Profiles of cytokines and cytotoxic proteins released by PBMCs were determined by the multiplex cytokine assay. The supernatant was collected at the indicated time points. Results were from three donors. MFI: mean fluoresce intensity. **B)** Viability of PBMCs on day 3. PBMCs cultured alone (w/o gel) were used to access spontaneous apoptosis. On day 3, PBMCs were stained with propidium iodide (PI) and PI positive PBMCs were considered as apoptotic/dead cells. Results were from three donors.

Next, the release of cytokines or cytotoxic proteins on days 1, 2 and 3 was analyzed. Out of the 13 examined cytokines/cytotoxic proteins, mainly IL-6, IFNγ and GzmB were detected in the medium. In *E. coli* gels, the release of IL-6 peaked on day 2 (Donor #1 and Donor #2) or day 1 (Donor #3) and reduced afterwards, whereas in *ClearColi* gels, peak days for IL-6 release vary among donors (day 3 for Donor #1, day 2 for Donor #2, day 1 for Donor #3) (Figure 5B). The peak level of released IL-6 was comparable (Donor #1) or lower (Donor #2 and Donor #3) in *ClearColi-* than in *E. coli*-gels (Figure 5B). Release of IFNγ in *ClearColi-*gels was detectable from day 1, which was not always the case for GzmB (Figure 5B). The peak levels of IFNγ or GzmB were comparable between *E. coli-* and *ClearColi*-gels (Figure 5B).

Considering that the number of viable immune cells in contact with *ClearColi*-gels was 1.5-2 fold higher than that with *E. coli*-gels, these results indicate that the release of inflammatory cytokines induced by the *ClearColi*-gels is considerably lower compared to *E. coli*-gels. The NK or T cell subsets on day 3 were analyzed via flow cytometry. No significant difference in the differentiation of NK cells, CD4^+^ T cells and CD8^+^ T cells was observed in *E. coli*- or *ClearColi*-gels compared to blank hydrogels (Figures 6A-F). Together, our findings indicate that physical separation of *ClearColi* or *E. coli* from immune cells by encapsulation in Pluronic hydrogels significantly reduces activation and changes in the differentiation of NK cells or T cells and, therefore, in the overall immune response.

**Figure 6:**
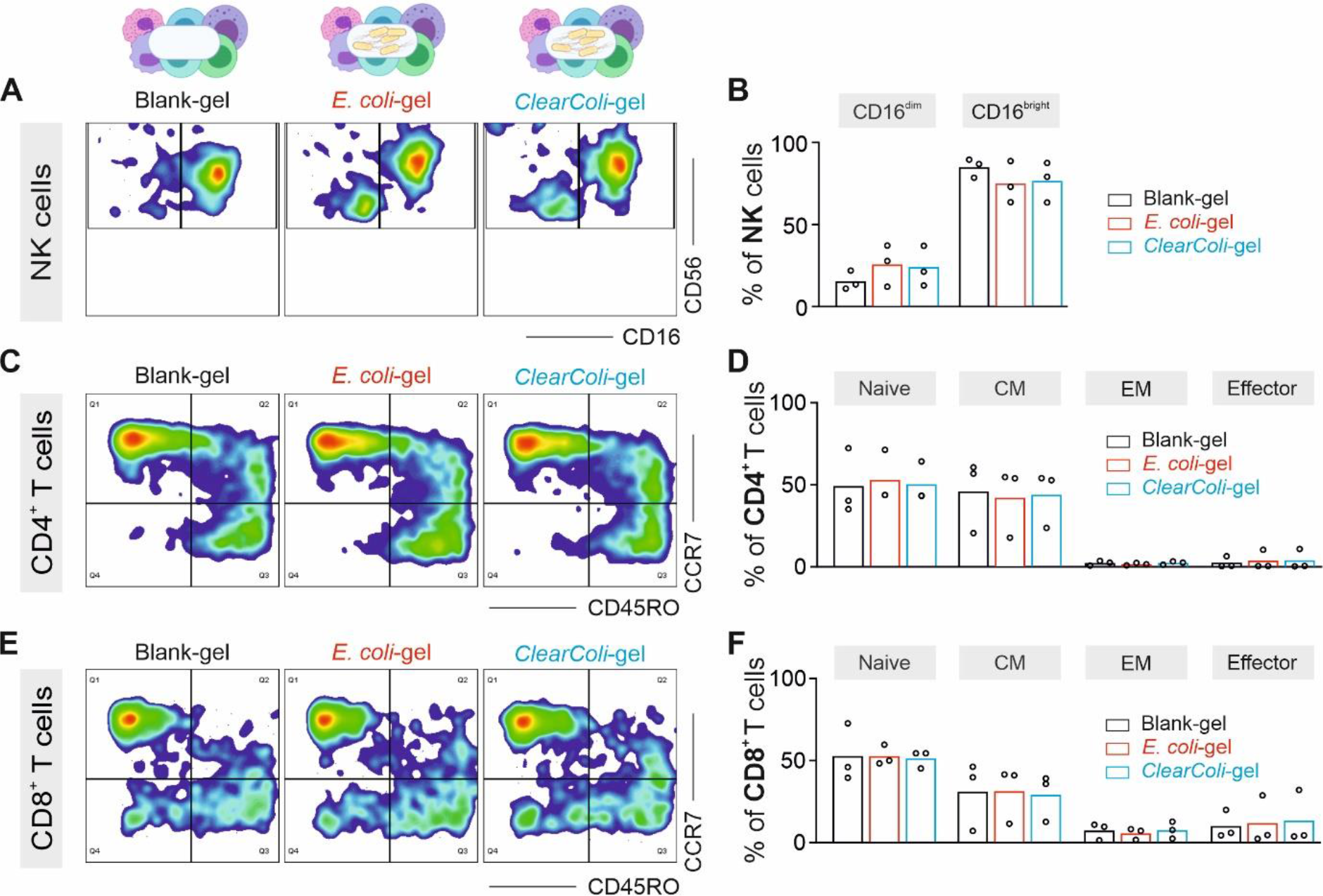
Differentiation of NK and T cells in response to bacteria encapsulated hydrogels. PBMCs were cultured with empty hydrogels (Blank-gel), *E.coli*-encapsulated (*E.coli*-gel) or *ClearColi*-encapsulated gels (*ClearColi*-gel). Differentiation of NK cells (**A, B**), CD4^+^ T cells (**C, D**) and CD8^+^ T cells (**E, F**) were examined using indicated surface markers. Results were from three donors. CM: central memory cells. EM: effector memory cells.

### Strong IFNγ release is induced in PBMCs from pre-activated donors

During the analysis, we noticed that out of six donors, PBMCs from three donors showed high spontaneous IL-2 release in the negative control (no hydrogel or bacteria) right from day 1 (Supplementary information Figure S5). Since IL-2 is mainly produced by activated T cells, especially CD4^+^ T helper cells, immune cells in these donors were very likely in a pro-inflammatory status (termed pro-inflammatory donors henceforth). For these donors, direct contact of PBMCs with bacteria induced large IFNγ release, with variable peak time among donors, and with similar peak levels for *E. coli* and *ClearColi* hydrogels (Figure 7A). IFNγ release was 3-30 times higher than in normal PBMCs (Figure 7B), whereas IL-6 release stayed in a comparable range (Figure 7C). GzmA was released in large quantities (Figure 7A) and about 10-200 fold higher than normal donors (Figure 7D). In the control experiments with non-encapsulated bacteria strong cytokine release was also observed (Supplementary information Figure S6).

**Figure 7:**
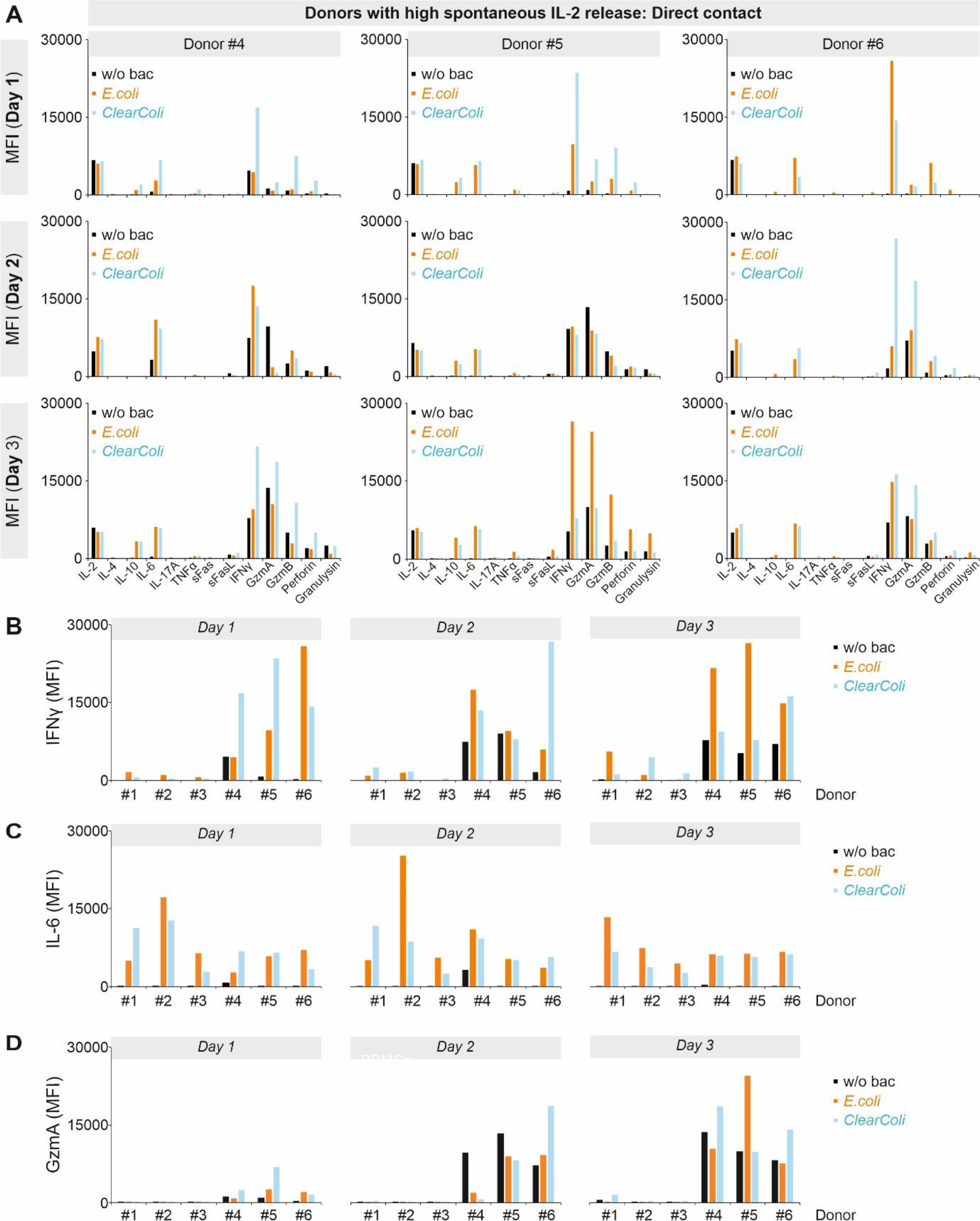
Profiles of cytokines and cytotoxic proteins from donors with high spontaneous IL-2 release upon bacterial contact. **A)** Profiles of cytokines and cytotoxic proteins released by PBMCs were determined by the multiplex cytokine assay. PBMCs with high spontaneous IL-2 release were cultured without bacteria (w/o bac) or in direct contact with *E.coli* or *ClearColi*. The supernatant was collected at the indicated time points. Results were from three donors. MFI: mean fluoresce intensity. **B-D)** Comparison of released cytokines between donors with low and high spontaneous IL-2 release. IFNγ, IL-6 and granzyme A (GzmA) are shown in **B, C** and **D**.

These pro-inflammatory PBMCs also showed prominent release of IFNγ, IL-6, GzmA, GzmB, perforin, and granulysin with both the blank and bacteria-containing gels (Figure 8, 9A). With the bacterial gels, IFNγ was 5-20 fold higher than in normal donors (Donors #1 and #2, Figure 9B). In comparison, levels of IL-6 were similar as the case of normal donors (Figure 9C). GzmA release was about 20-200 fold higher than normal donors (Figure 9A, D). Release of GzmB, perforin and granulysin were elicited in a considerably low level compared to IFNγ (Figure 9A). These results indicate that in pro-inflammatory donors featured with high spontaneous release of IL-2, *E. coli* and *ClearColi* LTMs could trigger considerable immune reactions.

**Figure 8:**
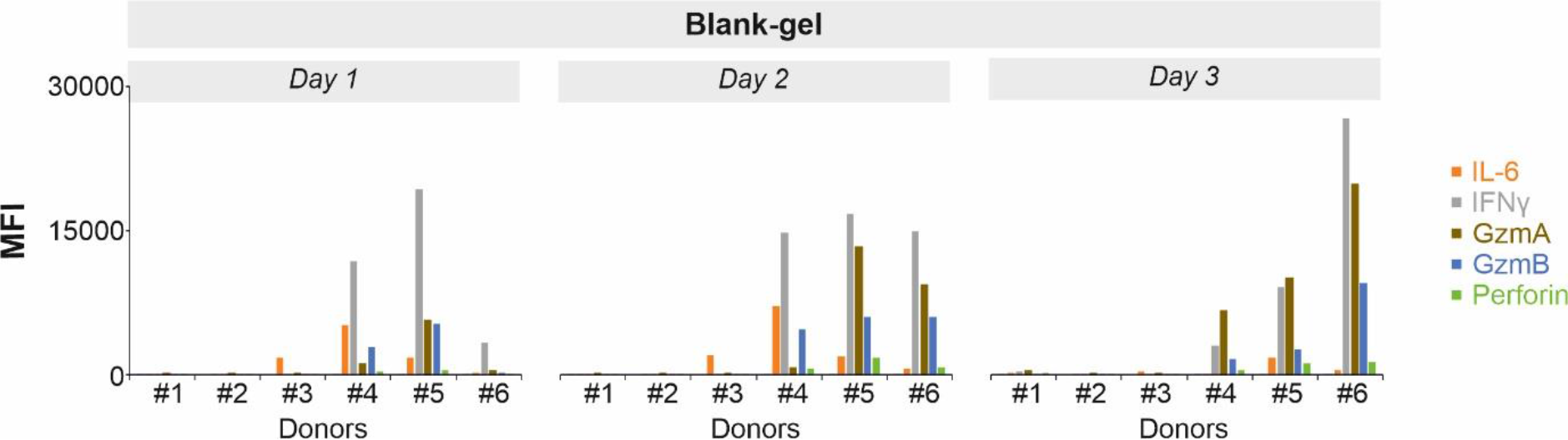
Comparison of cytokine profiles between donors with low or high spontaneous IL-2 release in response to empty hydrogels. PBMCs from Donors #1-3 were featured with low spontaneous IL-2 release, those from Donors #4-6) with high spontaneous IL-2 release. PBMCs were cultured with empty hydrogels (Blank-gel). The supernatant was collected at the indicated time points.IL-6, IFNγ, granzyme A (GzmA), granzyme B (GzmB), and perforin are shown. Data were extracted from Fig. 5A and Fig. 8A.

**Figure 9:**
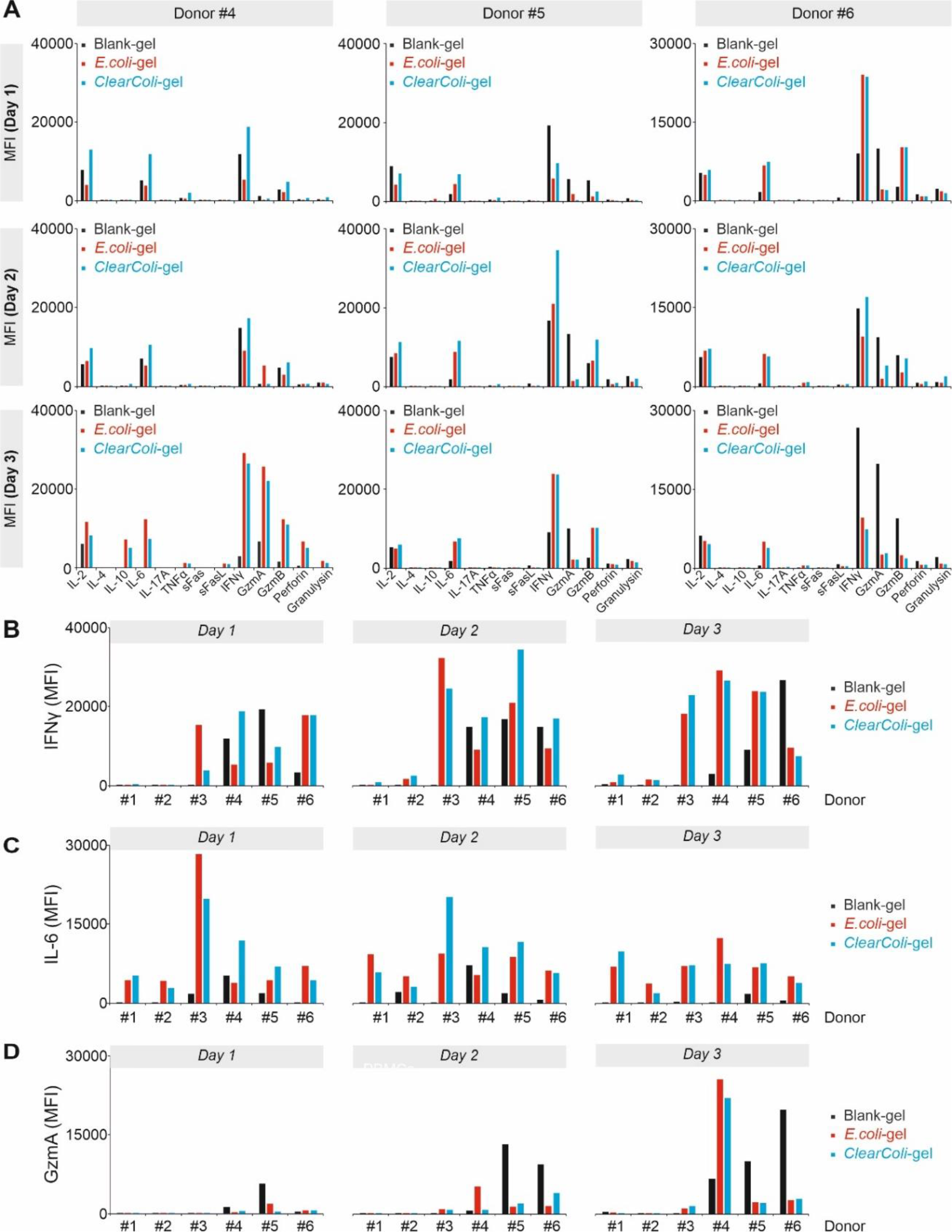
Profiles of cytokines and cytotoxic proteins from donors with high spontaneous IL-2 release in response to bacteria encapsulated hydrogels. **A)** Profiles of cytokines and cytotoxic proteins released by PBMCs were determined by the multiplex cytokine assay. PBMCs with high spontaneous IL-2 release were cultured with empty hydrogels (Blank-gel), *E.coli*-encapsulated (*E.coli*-gel) or *ClearColi*-encapsulated gels (*ClearColi*-gel). The supernatant was collected at the indicated time points. Results were from three donors, which are the same donors in Fig. 7. MFI: mean fluoresce intensity. **B-D)** Comparison of released cytokines between donors with low (Donors #1-3) and high (Donors #4-6) spontaneous IL-2 release. IFNγ, IL-6 and granzyme A (GzmA) are shown in **B, C** and **D**.

## Discussion

LTMs in direct contact with tissues and biological fluids could potentially trigger innate and adaptive immune responses in the body. These can be originated by biochemical (release of proteins, oligonucleotides, chemical moieties, etc.) or physical (mechanical properties of the construct) factors ^10, 46, 47^. Such responses have been observed during the development of encapsulated cell technologies, wherein mammalian cells engineered to secrete drugs are encapsulated in semipermeable matrices ^48^. Most studies were conducted in cells encapsulated in degradable alginate microcapsules, and immune responses were observed in terms of the polymer, the cellular antigens, the therapeutic transgenes and the DNA vectors used to genetically engineer the cells^48^. Improved designs can solve many of these issues by using semi-permeable membrane interfaces to physically separate the engineered cells from the host ^49^. On the other hand, engineered bacteria are being developed as live biotherapeutic products to be freely administered in the body for curing a variety of diseases like inflammatory bowels disease, viral infections, cancer, etc ^50, 51^. One of the prominent strains used in such studies is *E. coli* Nissle 1917, which is a probiotic in European markets. This strain has a defect in its LPS biosynthesis leading to truncated oligosaccharide chains that triggers low levels of immunogenicity when the LPS was exposed to PBMCs ^52^. This strain is predominantly used to treat diseases in its natural habitat, the gut, but its use in other sites of the body to treat cancer has also been explored ^53^. This has revealed that its intratumoral injection in mice leads to considerable increase in IL-6 and TNFα levels, which is desirable for tumor immunotherapy ^54^. Thus, outside the gut, this probiotic strain has been found to illicit immune responses. This strain is also susceptible to rapid immune attack and clearance from the body, which has been temporarily reduced by programming it to generate a surface capsular polysaccharide that encapsulates the bacterial cell ^55^. Probiotic microencapsulation is a common strategy used to protect the bacteria from harsh factors within the body during administration ^56^. However, the effect of encapsulation on immune responses has not been systematically evaluated. In this study, we have studied the response of immune cells to LTMs made of the most common LTM bacteria, *E. coli,* and a synthetic, biocompatible, non-degradable polymer, Pluronic F127 diacrylate. The LTMs were designed with a Plu/PluDA core hydrogel containing the bacteria, enveloped in a thin PluDA hydrogel with higher crosslinking that prevents bacterial escape. The hydrogel envelope prevents physical contact between the bacteria and the immune cells in the culture.

It should be noted that in transwell inserts, i.e. within the culture medium, bacteria can freely grow during culture time and the resulting bacteria density to which cells are exposed might be orders of magnitude larger than bacteria number physically constrained within the hydrogels. Based on our previous study, the bacteria within the gels can grow to colony sizes with a conservative mean volume of 250 μm^3^ ^28^ corresponding to ~400 cells (*E. coli* volume ~0.6 μm^3^). Thus, in the 2 μL of the bacterial gel, the bacterial numbers starting from 8 × 10^3^ cells are expected to reach between 10^6^-10^7^ cells. In the transwell inserts, at saturation levels of growth (OD 2 – 5) that are expected to be reached before day 3 in the 100 μL volume, the bacterial numbers are expected to reach between 10^8^ – 10^9^ cells. In direct contact with the PBMCs, a similar range can be expected although variations can occur due to killing of the bacteria by the immune cells. Additionally, since the colonies have already been allowed to form before initiation of the experiments with PBMCs, it is likely that growth, metabolic activity and death of the bacteria within these colonies will be occurring at a considerably diminished rate compared to those growing freely in medium. So, apart from physical separation, lower immune responses due to encapsulation could be caused by these additional factors.

Endotoxic LPS from Gram-negative bacteria, including *E. coli*, can elicit strong immune responses ^57^. The *ClearColi,* with no LPS, has been shown not to trigger NFκB activity ^26, 58^. In this work, we found that *ClearColi-* encapsulated hydrogels did not induced any changes in differentiation of NK cells and T cells, but elicited some release of the pro-inflammatory cytokines IL-6 and IFNγ from normal PBMCs. In this case, the levels of IL-6 and IFNγ are in a comparable range as the empty hydrogel-induced release in pro-inflammatory PBMCs. As empty Pluronic gels is shown/proven clinically safe, these levels of IL-6 and IFNγ can be considered as response from a normal ‘foreign body reaction’^59, 60^.

Of note, this *in vitro* study portrays an extreme case with high numbers (3 million) of immune cells in contact with the bacterial gels from the start. The bacterial density within the gels is estimated to be in the range of 10^9^ – 10^10^ cfu/mL, which is equivalent to a bacterial culture at saturation levels of growth and the range in which probiotic formulations are made (e.g. Mutaflor containing *E. coli* Nissle 1917 at 2.5–25 × 10^9^ cfu/mL). Although the numbers of immune cells recruited to sites of inflammation or wounds have not been explicitly quantified, from the reported *in vivo* studies ^61, 62, 63^, we estimate the numbers are in the range of a few thousand to a few hundred of thousand. Thus, the measured response is very likely to be amplified/exaggerated compared to *in vivo* scenarios. Therefore, we conclude that *ClearColi*-encapsulating hydrogels do not elicit severe adverse events and are suitable for further *in vivo* studies.

Our results with PMBCs from pro-inflammatory donors indicate that LTMs might induce an elevated inflammatory reaction at the implantation site compared to donors with the immune system at a quiescent status. Considering the time of sample collection (between October and December 2020, a high season for seasonal flu and normal cold overlapped with the second wave of COVID-19 in Germany) the probability that some donors got infection a few weeks before the donation but were symptom-free at the time of donation was high. In this case, it is possible that the immune system could not have retracted to a quiescent status yet. The substantial difference between donors with low or high spontaneous IL-2 release highlights the necessity to perform detailed studies of the immune response to LTMs, including individuals with different immune status. Implementation of inflammation-controlling features ^64^ in LTMs, such as a temporal release of anti-inflammatory compounds from the material or the bacteria, could benefit recipients with a pro-inflammatory status.

## Author Contributions

AKY performed PBMC-related experiments and the corresponding analysis if not mentioned otherwise; SB prepared hydrogels, bacteria and bacteria-encapsulated hydrogels and carried out quality check; AdC helped with data interpretation and provided critical feedback on all aspects of the project; SS and BQ generated concepts; SS and SB designed hydrogel-related experiments and BQ designed PBMC-related experiments; All authors contributed to the writing, editing and cross-checking of the manuscript.

## Acknowledgments

We thank the Institute for Clinical Hemostaseology and Transfusion Medicine for providing donor blood; Markus Bischoff and Philipp Jung for the bacteria culture facility, Carmen Hässig, Cora Hoxha, and Gertrud Schäfer for excellent technical help; This project was funded by the Deutsche Forschungsgemeinschaft (SFB 1027 Project A2 to BQ, B6 to AdC, B8 to SS and GZ: INST 256/419-1 FUGG) and the Leibniz Science Campus on Living Therapeutic Materials, LifeMat.

## Ethical considerations

Research carried out for this study with material from healthy donors (leukocyte reduction system chambers from human blood donors) is authorized by the local ethic committee Ethik-Kommission bei der Ärztekammer des Saarlandes (Identification Nr. 84/15, Prof. Dr. Rettig-Stürmer).

