## Supplementary information for "*In vitro* evaluation of immune responses to bacterial hydrogels for the development of living therapeutic materials"

Bin Qu

Biophysics, Center for Integrative Physiology and Molecular Medicine (CIPMM)
School of Medicine, Saarland University, 66421 Homburg, Germany.
INM - Leibniz Institute for New Materials, 66123 Saarbrücken, Germany.


Shrikrishnan Sankaran

INM - Leibniz Institute for New Materials, Campus D2 2, 66123 Saarbrücken, Germany.

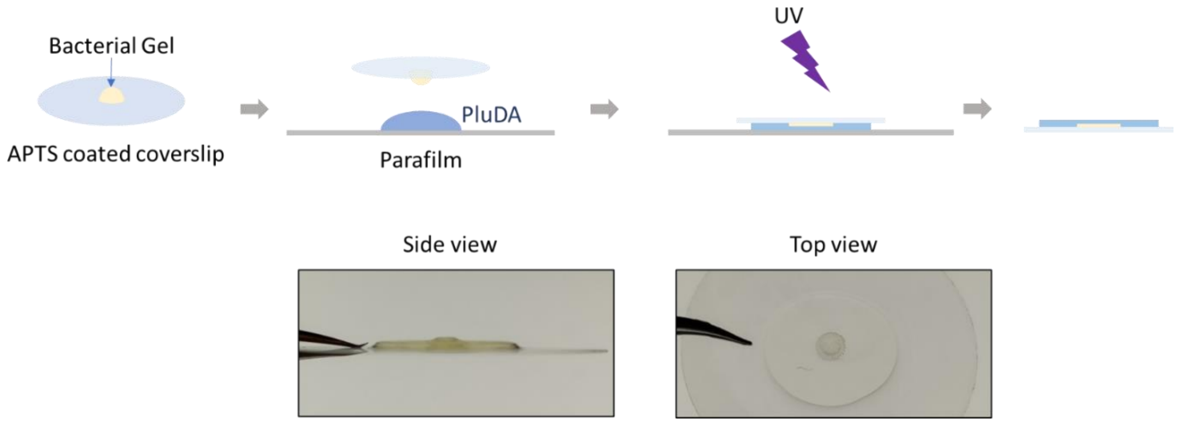


**Figure S1: Schematic of the fabrication of a leak-proof thin-film bacterial hydrogel construct.** The encapsulating gel, covalently crossliked to the APTS silanized glass coverslip upon UV irradiation, prevented the escape of bacteria from the bacterial gel upon contact with the surrounding medium.

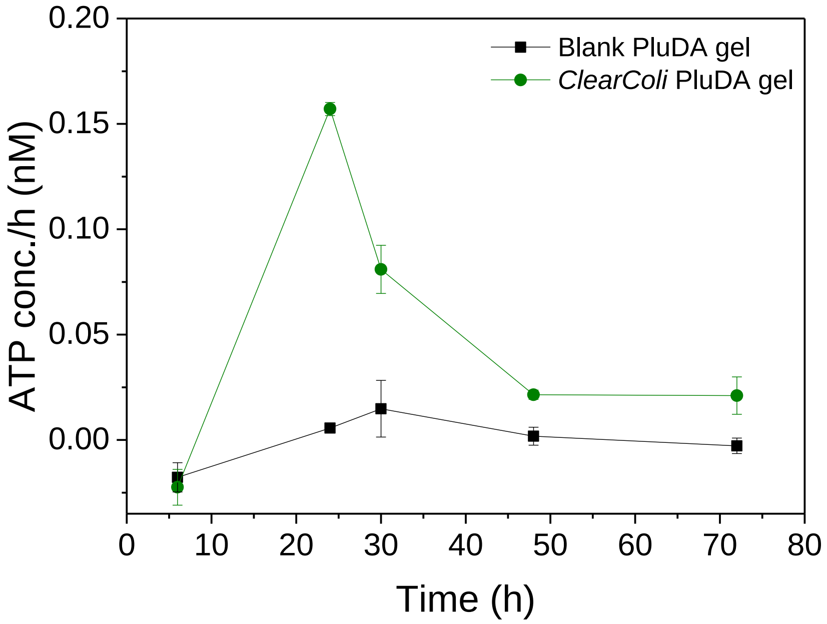


**Figure S2:** Extracellular ATP level changes in the surrounding medium: Samples were collected from the surrounding medium of bilayer thin films containing Clearcoli bacteria at timepoints 6, 24, 30, 48 and 72 h after fabrication, with blank bilayer thin films (no bacteria) as control. The concentration of the ATP was determined using a standard curve of ATP standard solution serially diluted in culture medium. (Symbols are means from 2 experimental replicates with 2 technical replicates each, error bars represent standard deviation)

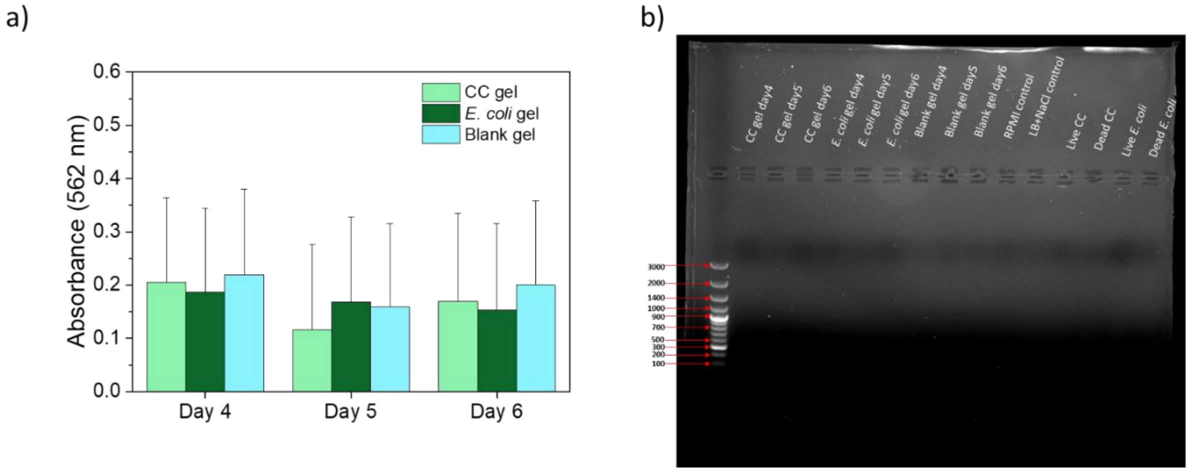


**Figure S3: Protein and DNA release by the bacteria in encapsulated gels. A)** The amount of protein released by the bacteria in the surrounding medium was measured using the BCA protein assay. The absorbance at 562 nm for the surrounding medium of the bacterial gels showed no significant difference compared to the blank gels; **B)** Agarose gels showing no indication DNA fragments released by the bacterial gels to the surrounding medium.

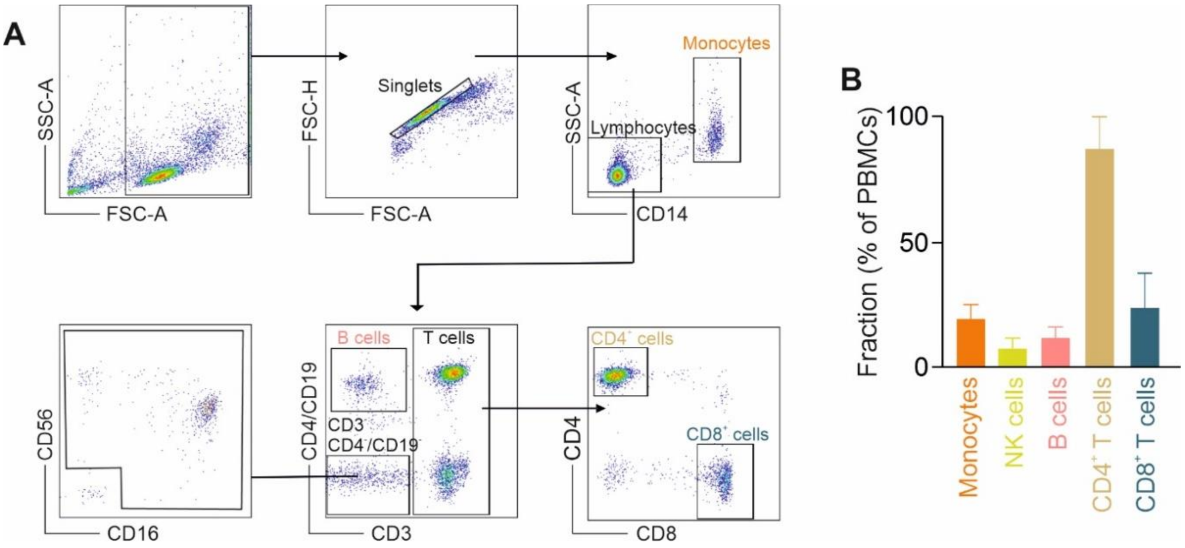


**Figure S4: Sub-populations of PBMCs from healthy donors. A)** Immunostaining was performed on PBMCs isolated from healthy donors and data was acquired using flow cytometer. Different immune cell population were gated from the lymphocytes as shown in the gating strategy by using the following antibodies with fluorophore conjugates: CD14-PE-Cy7, CD3-PerCP, CD4-BV421, CD19-BV421, CD56-APC, CD16-PE, and CD8-FITC as reported previously {Knorck, 2018 #66}. **B)** Fraction of each subpopulation in PBMCs. Results are from six donors.

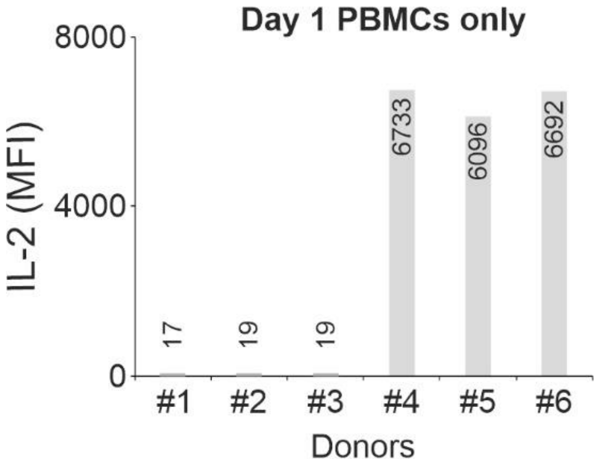


**Figure S5: Spontaneous release of IL-2 in different donors.** PBMCs from Donors #1-3 were featured with low spontaneous IL-2 release, those from Donors #4-6) with high spontaneous IL-2 release. PBMCs were cultured alone without bacteria or gels. The supernatant was collected on day 1. Data were extracted from Fig. 3 (Day 1) and Fig. 7A (Day 1).

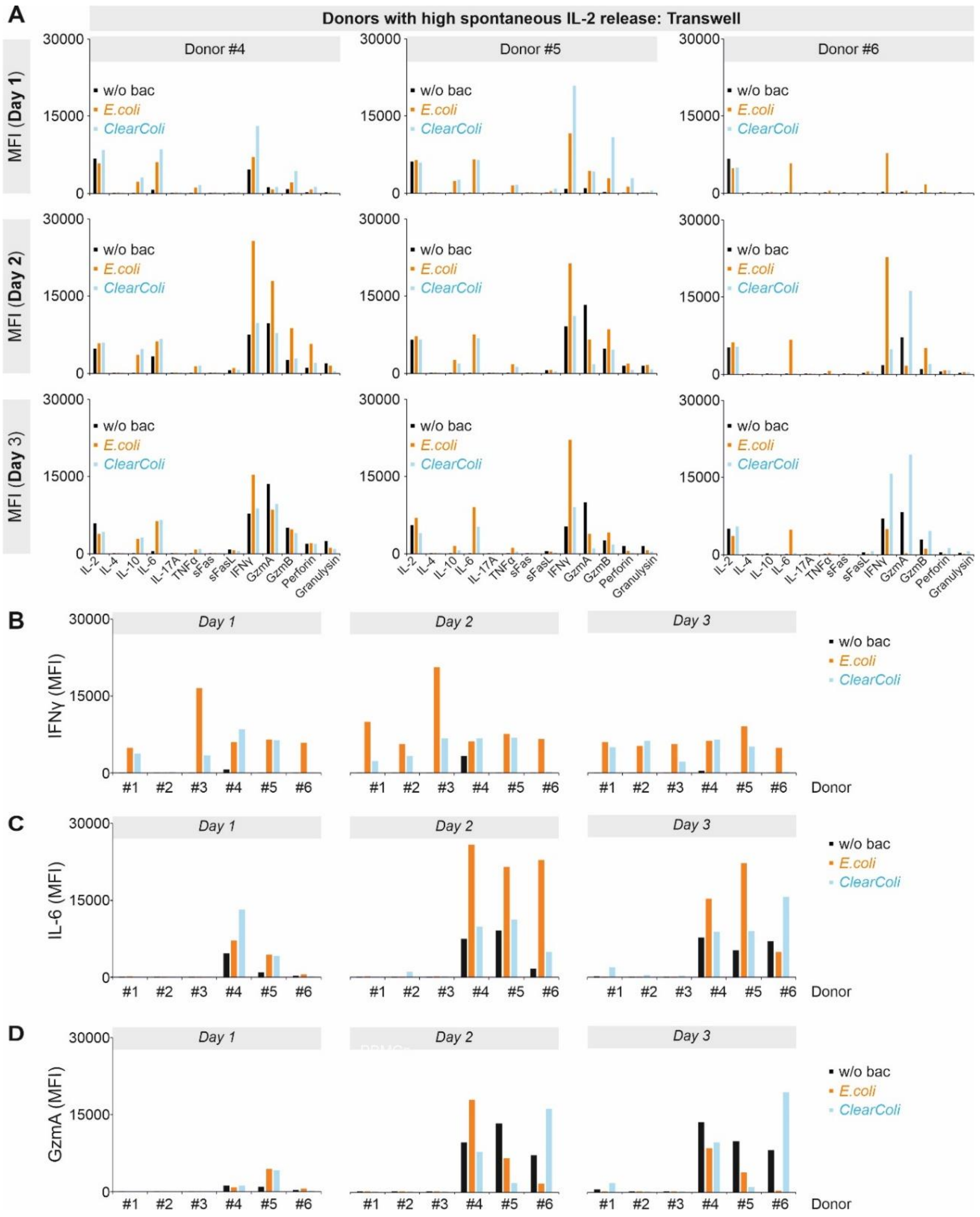


**Figure S6: Cytokine profiles of PBMCs from donors with high spontaneous release of IL-2 with E.coli or ClearColi bacterial strains separated by a nano-porous transwell insert. A**) PBMCs from donors with high spontaneous release of IL-2 were incubated with either *E.coli* or *ClearColi* strains separated by a nano-porous transwell insert for 3 days. Cell culture supernatant was collected on days 1, 2 and 3 to perform a multiplex cytokine assay. Cytokines and cytotoxic proteins released by PBMCs in response to soluble factors released by the bacteria was determined which was compared to w/o bacteria condition (PBMCs alone). The data is represented with Mean fluorescence intensity (MFI) on Y-axis for donors 4, 5 and 6 for all the 3 days (n=3). **B-D**) A comparison between donors with modest release of IL-2 (donors 1,2,3) and high spontaneous release of IL-2 (donors 4,5,6) for IFNγ (**B**), IL-6 (**C**) and Granzyme A (**D**) released from PBMCs incubated with *E. coli* or *ClearColi* strains in a transwell insert and w/o bacteria condition (PBMCs alone).
